# It’s not easy seeing green: the veridical perception of small spots

**DOI:** 10.1101/2022.11.15.516626

**Authors:** John Erik Vanston, Alexandra E. Boehm, William S. Tuten, Austin Roorda

**Author notes:** Bioengineering Research Partnership Grant, National Eye Institute: R01EY023591. Center Core Grant, National Eye Institute: P30EY003176. Training Grant, National Eye Institute: T32EY007043. Air Force Office of Scientific Research Awards FA9550-20-1-0195 & FA9550-21-1-0230. <u>**Disclosures:**</u>. **JEV:** none. **AEB:** none. **WST:** University of California (P). **AR:** University of California (P); University of Houston & University of Rochester (P).

## Abstract

When single cones are stimulated with spots of 543-nm light presented against a white background, subjects report percepts that vary between predominately red, white, and green. However, light of the same spectral composition viewed over a large field under normal viewing conditions looks invariably green and highly saturated. It remains unknown what stimulus parameters are most important for governing the color appearance in the transition between these two extreme cases. The current study varied the size, intensity and retinal motion of stimuli presented in an adaptive optics scanning laser ophthalmoscope. Stimuli were either stabilized on target locations or allowed to drift across the retina with the eye’s natural motion. Increasing both stimulus size and intensity led to higher likelihoods that monochromatic spots of light were perceived as green, while only higher intensities led to increases in perceived saturation. The data also show an interaction between size and intensity, suggesting that the balance between magnocellular and parvocellular activation may be critical factors for color perception.

Surprisingly, under the range of conditions tested, color appearance did not depend on whether stimuli were stabilized or not. Sequential activation of many cones does not appear to drive hue and saturation perception as effectively as simultaneous activation of many cones.

## Introduction

Color perception depends on the activity of three types of cone photoreceptors. Each individual cone is unable to distinguish between light intensity and spectral composition, which is called the principle of univariance (Rushton, 1972). Therefore, color vision is, at the microscopic scale, a spatial task, where the signals of the three cone types are compared across discrete units of retinal space. Because of this, a brief, small and dim spot of narrowband light designed to stimulate single photoreceptors delivered to the retina will vary in color appearance as the light stimulates different retinal locations, as demonstrated by (Krauskopf & Srebro, 1965). Hofer, Singer, and Williams (2005) performed a similar experiment, delivering small, brief spots of narrowband light to the retina, with significantly improved optical confinement of the spot through the use of adaptive optics to correct for blur from ocular aberrations. They found that hue percepts elicited by single cone sized stimuli are highly variable when viewed against a dark background. In neither of these studies were the investigators able to target specific cone types.

More recent studies used image-based eye tracking (Arathorn et al., 2007; Harmening, Tuten, Roorda, & Sincich, 2014; Yang, Arathorn, Tiruveedhula, Vogel, & Roorda, 2010) in an adaptive optics scanning laser ophthalmoscope (AOSLO) system to accurately target and deliver stimuli to single cones and investigate the percepts elicited thereby. Presenting 543-nm stimuli on a white background that equated the baseline adaptive state of the three cone types in eyes whose cone mosaics had been mapped, Sabesan, Schmidt, Tuten, and Roorda (2016) and Schmidt, Boehm, Foote, and Roorda (2018) showed that the spectral identity of each cone is recapitulated in its color percepts. While many stimuli appeared white, monochromatic light delivered to L cones was more likely to appear red, and the same light delivered to M cones was more likely to appear green. Moreover, repeated stimulation of the same cone produced consistent color sensations. Expanding on this paradigm to test pairs of cones simultaneously, Schmidt, Boehm, Tuten, and Roorda (2019) showed that when pairs of cones are stimulated, the resulting hue percept was predicted by the average of the hue percepts of each cone when stimulated individually. They further showed that when like pairs of cones (e.g., two L cones) were stimulated, the resulting percept was more saturated than that elicited by either cone on its own. When an L cone and an M cone were stimulated at the same time, the resulting percept was less saturated than when either cone was stimulated in isolation.

However, narrowband 543-nm light viewed over a large field and under normal viewing conditions looks invariably green and highly saturated. There are several important differences between the conditions in the studies cited above and normal viewing. One difference is the sheer number of cones engaged; even a 2-degree spot, considered to be rather small in vision science, will deliver light to as many as 20,000 cones when viewed foveally (Curcio, Sloan, Kalina, & Hendrickson, 1990; Wang et al., 2019).

In the transition between these two extremes, several stimulation parameters govern the number of cones that sample the stimulus: blur, size and motion. Blur and size are easily controlled in the AOSLO and are fully expected to have effects on color appearance. The role of motion, on the other hand, is less predictable. Small eye movements are ubiquitous in natural vision, and are leveraged to increase contrast at high spatial frequencies (Rucci, Iovin, Poletti, & Santini, 2007) and to improve fine spatial vision (Anderson, Ratnam, Roorda, & Olshausen, 2020; Ratnam, Domdei, Harmening, & Roorda, 2017). Are small eye movements leveraged in a similar way to build a more veridical color percept?

It remains unknown the extent to which each of these stimulus parameters are used in determining the color appearance of very small spots. In the current study we investigated this issue, aiming to answer the following questions: how many cones must be stimulated to achieve a veridical (in this case, green) color percept? Does it matter if this is achieved through increasing the stimulus size, or by allowing a small stimulus to move across the retina? And finally, while previous research has shown that varying the intensity of single-cone stimuli does not affect hue perception (Schmidt et al., 2018), does it have an effect for larger stimuli?

## Methods

### Subjects

Four experienced subjects participated in these experiments (median age 33, N = 2 female). All subjects had normal color vision, and two (S20210 & S10003) were authors of this study. All procedures were approved by the Institutional Review Board at the University of California Berkeley and adhered to the tenets of the Declaration of Helsinki. Informed consent was obtained from each subject before the experiments. At the start of each session, cycloplegia and mydriasis were induced with drops of 1.0% tropicamide and 2.5% phenylephrine hydrochloride ophthalmic solution.

### AOSLO microstimulator

An adaptive optics scanning laser ophthalmoscope (AOSLO) was used for retinal imaging and stimulus presentation. The AOSLO system has been described in detail elsewhere (Mozaffari, LaRocca, Jaedicke, Tiruveedhula, & Roorda, 2020; Roorda et al., 2002; Yang et al., 2010). Briefly, light from a supercontinuum laser (SuperK Extreme; NKT Photonics, Birkerod, Denmark) was split into three independent channels with central wavelengths of 940, 840 and 543 nm. The light from each channel was launched into the system via a single mode fiber. The relative vergence of each channel was adjusted to be equal and opposite to the average longitudinal chromatic aberration of the human eye (Atchison & Smith, 2005; Grieve, Tiruveedhula, Zhang, & Roorda, 2006) and they were coaligned and directed into the AOSLO system. All beams were focused on the retina and scanned in a raster pattern using a fast 16 kHz resonant scanner (Electro-Optical Products Corp, Fresh Meadows, NY) and a galvanometric scanner scanning a sawtooth at 30 Hz.

Returned light from the 940-nm channel was sent to a wavefront sensor that measured optical aberrations, which were continuously corrected by a deformable mirror (DM97-08; ALPAO, Montbonnot-Saint-Martin, France) in a closed-loop manner.

Returned light from the 840-nm channel was sent through a confocal pinhole and collected by a photomultiplier tube (H7422-50; Hamamatsu, Hamamatsu City, Japan) and used to generate a 512x512 pixel, 30-frame-per-second, live video of the retina over a 0.9x0.9 degree field. The live video was also used to track the retina using strip-based cross-correlation as described in (Arathorn et al., 2007).

The 543-nm light was used to stimulate the retina. 543 nm was chosen because L and M cones have approximately equal sensitivity to this light, and S cones are insensitive to it. High speed acousto-optic modulators (Brimrose Corp, Glencoe, MD) controlled the 543-nm power at precise time points during the raster scan to project spatial patterns to the retina. The exact location was guided by the real-time tracking. Transverse chromatic aberration (TCA) between the 840 and 543-nm wavelengths was measured and corrected during each session using the image-based method described in Harmening, Tiruveedhula, Roorda, and Sincich (2012). Since TCA is highly dependent on the position of the AOSLO beam in the pupil, two measures were taken to control it. First, the subject used a dental impression mount fixed to an X-Y-Z stage to minimize eye movement during the experiment. Second, a CCD camera was integrated into the system to view a retro-illumination image of the subject’s pupil using a fraction of the 940-nm wavefront sensor light. The exact eye position was marked at the time TCA was measured and was continuously monitored during the experiment. The experimenter adjusted the X-Y alignment of the eye to maintain a constant position via tracking of the first Purkinje reflex (Domdei, Linden, Reiniger, Holz, & Harmening, 2019).

### Retinal areas

Previous studies have examined the repeatability of hue judgments for stimuli delivered to individual cones and pairs of cones (Sabesan et al., 2016; Schmidt et al., 2018; Schmidt et al., 2019). In the current study, a different group of 6-7 target locations centered on cones were chosen on each session. These locations were chosen to be between 0.5- and 1-degree eccentricity either nasal or temporal to the fovea. During each session a high-quality reference image was created by averaging together 60-150 frames of video from the region of interest. Target locations in this study were chosen from these reference images.

### Background & fixation

An external projector (DLP4500; Texas Instruments, Dallas, TX) was used to provide a background and a fixation spot. The projector was integrated into a Badal optometer to correct for each eye’s spherical refractive error. A beam splitter placed in front of the subject’s eye was used to integrate the image from the projector into the path of the AOSLO beam so that at any time the laser raster and the field of the projector were superimposed. At the start of each experimental session subjects co-registered the projector and AOSLO raster by using arrow keys on a keyboard to align a white cross viewed on the projector with the center of the AOSLO raster. A small fixation spot was then placed relative to that cross at the desired eccentricity, and the raster-alignment cross was turned off.

Once this alignment procedure was complete, the entire field of the projector was filled with white light metameric with equal energy white; this provided a background for the experiment and ensured that all three cone types had roughly equal levels of excitation in their adapted state. Subjects used a keyboard to adjust the intensity of this white background until the 840-nm raster was just barely visible. The mean luminance across sessions was 9.92 cd/m^2^ (standard error of the mean = 1.82). The fixation target was a black dot that appeared as a decrement against this background. All procedures described above were drawn using the Psychophysics Toolbox extensions (Brainard, 1997; Kleiner et al., 2007; Pelli, 1997) for MATLAB (The Mathworks, Natick, MA).

### Display characterization

Light from the projector entered the eye in Maxwellian view, complicating direct radiometric characterization of its LED primaries. To measure their spectral power distributions (SPDs), we used a combination of 1) direct measurements of the projector’s primary spectra at many intensity levels using a PR-650 spectroradiometer (Photo Research, Los Angeles, CA), with the objective lens of the PR-650 at the location of the subject’s eye, and 2) the method of psychophysically equating Maxwellian and Newtonian view luminances described in Leibowitz (1954). A bipartite field was delivered to a subject’s dominant eye, with one half of the field delivered by the projector via Maxwellian view, and the other half delivered by a LCD monitor via Newtonian view. The intensity of the Newtonian half field was set to maximum, and the Maxwellian half of the field was adjusted in intensity until the two halves of the field appeared equally bright. This brightness matching task was repeated 10 times and the settings averaged. The luminance of the average match brightness was then measured in Newtonian view using the PR-650. These measurements were done using the blue primary from both displays, as their spectra were similar enough that at equiluminance, the bipartite field appeared perfectly uniform.

### Measurement of inter-cone distances

Stimulus sizes used in the current study were defined by the distances between cone centers in the region of the retina being tested on the day of a given experimental session. These inter-cone distances were defined based on the reference image (described above in “Retinal areas”). Because residual local image distortions due to eye motion tend to occur at the edges of the stabilized image, only the center 50% of the reference image was considered. From this region, 100 “seed cones” were selected randomly without replacement. For each seed cone, the distances between its center and the centers of all other cones in the region were calculated. The six nearest cones were defined as its nearest neighbors, and the twelve nearest cones beyond the nearest neighbors were defined as its second nearest neighbors. Due to the generally hexagonal packing of cone photoreceptors at this eccentricity, the nearest neighbors were likely to be in a ring directly surrounding the seed cone in question, and the second nearest neighbors were likely to be in an approximate ring surrounding the nearest neighbors. This process was repeated for each seed cone, yielding 600 nearest neighbor distances (NND) and 1200 second nearest neighbor distances (SNND); these values were averaged within each group to compute a mean NND and a mean SNND. For this study, one stimulus size was intended to cover a target cone and its surrounding cones, and another stimulus size was intended to cover a target cone and two concentric rings of cones. To achieve these coverages, the mean NND and SNND were doubled and 3 pixels (0.32 arcmin) were added to encompass the entire cone. Median stimulus sizes were 0.54 arcmin, 2.3 arcmin, and 3.97 arcmin.

### Detection thresholds

Stimulus intensities used in the current study were defined by multiples of detection thresholds at each stimulus size. On each session, 6-7 target locations centered on cones were chosen from the reference image. On each trial, subjects used keypresses to trigger stimulus onsets and respond whether they saw the stimulus or not.

The stimulus comprised a circular spot of 543-nm light delivered to one of the target locations, stabilized on the retina and varied pseudorandomly across trials. Stimulus intensity was tracked with two randomly interleaved QUEST staircases (Watson & Pelli, 1983) that aimed for 85% frequency of seeing and terminated after 30 trials per staircase. The threshold estimates from the two staircases were then averaged together to yield a detection threshold for that stimulus size and for that session. Detection thresholds for each stimulus size were measured separately. The method used here, in which different target locations are stimulated across trials within the same staircases, yields an average threshold across a region of the retina.

The experimenter watched a live video stream of the subject’s retinal mosaic, wherein the light delivery location was indicated by a digital marker onscreen. Trials that were determined by the experimenter to contain stimulus misdeliveries were repeated.

### Hue scaling

Stimuli in the hue scaling task were spots of 543-nm light delivered to the same target locations as in the detection threshold task for each session. Trial duration was 0.5 seconds, comprising 15 brief deliveries over 15 AOSLO raster frames.

Three stimulus sizes were used, designed to stimulate single cones (“small” size, 0.54 arcmin), a target cone and the surrounding six cones (“medium” size, 2.3 arcmin), and a target cone and two surrounding concentric rings of cones (“large” size, 3.97 arcmin, encompassing ∼19 cones).

For the medium and large sizes, two stimulus intensities were shown: one at detection threshold and the other at either six times that threshold or the maximum intensity of the laser with the chosen neutral density filter, whichever was lower. Previous studies have shown that increasing the intensity of monochromatic light delivered to single cones does not affect the resulting hue or saturation (Schmidt et al., 2018), so small stimuli in the current study were only shown at threshold intensity.

To determine the impact of eye motion on hue and saturation perception, two eye motion conditions were used. In the “stabilized” condition, stimuli were delivered to their targets using the same stabilization method used in the detection threshold task. As the eye moved, its motion was tracked against a reference frame using strip-based cross-correlation, and the stimulus was moved commensurately so that the light was delivered to a single location on the retina over the course of the trial (Arathorn et al., 2007). In the “unstabilized” condition, stimuli were initially delivered to the same target locations as in the stabilized condition, but after the first stimulus frame (33 ms) stabilization was turned off, allowing the stimulus to be sampled freely by the moving photoreceptor mosaic. As a result, the total energy delivered was the same, but the number of cones stimulated on a given trial in the unstabilized condition was higher than in the stabilized condition.

On each trial, subjects pressed a key to trigger a stimulus presentation, repeating the presentation if prompted by the experimenter. Subjects used keypresses to rate the saturation of the stimulus with an integer from 1 to 5, where 1 was the least saturated and 5 was the most saturated. They then used keypresses to describe the hue of the stimulus: was it more greenish, more reddish, or pure white? On trials where the subject described the stimulus as pure white, the saturation was retroactively coded as zero. If the stimulus could not be perceived by the subject on that trial, they could indicate “not seen”, which moved to the next trial without any further responses. Since previous research using cone-sized stimuli presented on a white background showed that the majority of the variation in hue responses occurs along a red-green axis that passes through white (Schmidt et al., 2018; Schmidt et al., 2019), we considered these hue responses to be sufficient to detect systematic variation in the dependent variable.

A given experiment session included two stimulus sizes (small & medium or medium & large), two intensities, two eye motion conditions, and 6-7 target locations, pseudorandomly interleaved. The number of trials per session was limited by the subject’s fatigue and by the duration of mydriasis & cycloplegia; 150-250 trials were typically completed per session.

### Trial exclusion

For every trial, a video was saved of the subject’s retina, with an overlaid digital marker indicating the exact location of the stimulus on each frame. These stimulus locations were extracted from each video for further analysis. Videos were 1 second in duration with a framerate of 30 Hz; the stimulus was on for 15 frames within each 1-second video.

Trials were selected for inclusion in the final analysis based on how successfully the stimulus was delivered to its intended target. Different trial exclusion criteria were used for the two eye motion conditions since the intended behavior of the stimulus differed between the two. Both conditions’ exclusion criteria were based on the statistics of the observed deliveries.

For the stabilized condition, it was intended for the stimulus to be consistently delivered to the target location despite the movement of the eye. For each trial, the distance between the target pixel and the delivery pixel (the pixel that the center of the stimulus was actually delivered to) was calculated for each frame. For frames on which the target pixel and the delivery pixel were different, the direction of the delivery error was not of interest; therefore, only the absolute distance was considered. Because these distances cannot be less than zero, distributions of delivery errors were positively skewed, and so the median was used as a measure of central tendency and interquartile ranges (IQR) were used as a measure of variation. A median delivery error was computed for each trial, and these trial median errors were considered across all sessions for all subjects when determining the threshold for excluding individual trials. The grand median delivery error for stabilized trials was 2.24 pixels (∼0.24 arcmin). Stabilized trials were excluded if their median delivery error was greater than 1.5 times the IQR above the median (3.64 pixels, or ∼0.39 arcmin). 242 out of 1793 stabilized trials were excluded in this way (13.50%).

For the unstabilized condition, the stimulus was delivered to the target location on the first frame, and then stabilization was turned off and the stimulus was allowed to drift across the retina due to natural fixational eye motion. For this condition the distance between the target pixel and the delivery pixel was only relevant for one frame, and thus this first-frame delivery error (FFDE) was used as the exclusion criterion. The grand median FFDE for unstabilized trials was 2.24 pixels (∼0.24 arcmin). Unstabilized trials were excluded if their FFDE was greater than 1.5 times the IQR above the median (6.30 pixels, or ∼0.68 arcmin). 272 out of 1813 unstabilized trials were excluded in this way (15%).

Trials wherein the subject reported not seeing the stimulus despite an accurate delivery were also excluded from further analysis. 558 trials out of a total of 3606 (15.47%) met this exclusion criterion. Considering all three of these criteria, 2594 trials were included in the final data analysis.

### Data analysis

The two dependent variables in this study were saturation rating and hue name. For each subject the saturation ratings for each condition were averaged across trials. Because there are no numerical values associated with hue names, the proportion of trials that each hue response was given was considered for each condition and for each subject. These proportions were then averaged across subjects when different conditions were compared. Since 543-nm light looks invariably green under normal viewing conditions, the proportion of “green” responses were singled out for analysis.

The data in this study are based on a fairly large number of trials from a small number of subjects; additionally, due to practical considerations, one subject only completed the medium and large conditions. While parametric testing with small sample sizes is not out of the question, the unequal sample sizes across conditions motivated us to use nonparametric tests. Statistical comparisons were made using the Kruskal-Wallis test, which can be considered a non-parametric version of the one-way ANOVA (Kruskal & Wallis, 1952). However, the conclusions that can be drawn from these tests are limited by low statistical power, which makes it relatively more likely that a Type II error will be committed.

## Results

In this section, each pair of independent variables will be considered in turn. Figure 1 shows the mean saturation ratings and proportion green responses for all subjects, collapsed across stimulus intensities to examine stimulus size. There was no significant effect of stimulus size on mean saturation ratings [χ^2^(2) = 0.84, p = 0.66], nor was there a significant effect of eye motion condition on mean saturation [χ^2^(1) = 0.57, p = 0.45]. There was a significant effect of size on the mean proportion of green responses [χ^2^(2) = 14.57, p = 0.001, η^2^ = 0.66], with larger stimuli being more commonly reported as appearing green. Eye motion condition had no significant effect on the proportion of green responses [χ^2^(1) = 0.01, p = 0.92].

**Figure 1:**
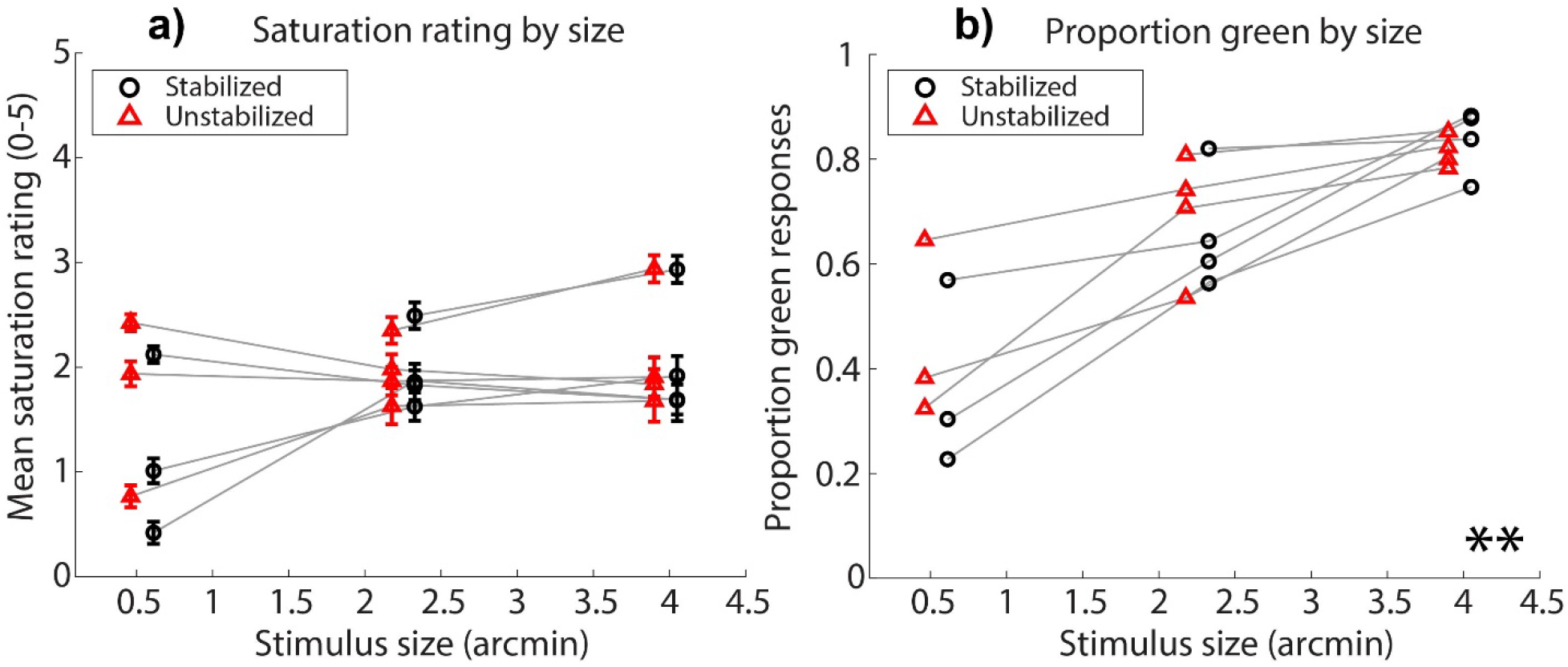
**a)** Saturation rating for each stimulus size condition. Circles are mean saturation ratings for each subject across trials for stabilized stimuli; triangles are for unstabilized stimuli. Error bars represent +/-1 standard error of the mean across trials. **b)** Proportion of trials that subjects described stimuli of each size as green, as opposed to red or white. Circles are green proportions for stabilized stimuli; triangles are for unstabilized stimuli. ** indicates that there was a main effect of size where p < 0.01. In both panels, data are collapsed across intensity conditions, and horizontally offset from the true abscissa value for aesthetic purposes.

Figure 2 shows the mean saturation ratings and proportion of green responses for all subjects, collapsed across stimulus sizes to examine stimulus intensity. Increasing stimulus intensity was effective at increasing both mean saturation ratings [χ^2^(1) = 6.89, p = 0.01, η^2^ = 0.42] and proportion of green responses [χ^2^(1) = 11.29, p = 0.001, η^2^ = 0.74]. Eye motion condition had no significant effect on either saturation [χ^2^(1) = 0.01, p = 0.92] or proportion green [χ^2^(1) = 0.18, p = 0.67].

**Figure 2:**
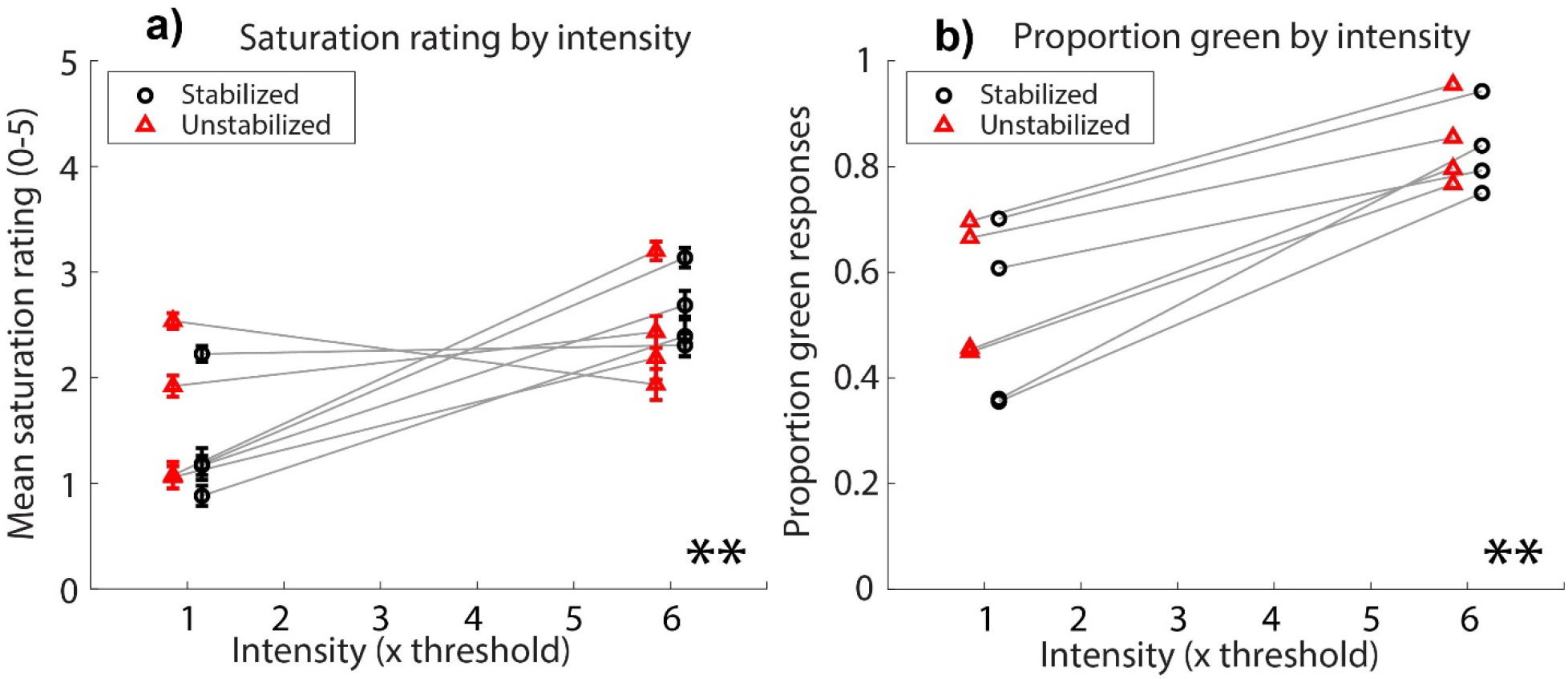
**a)** Saturation rating for each stimulus intensity condition. Circles are mean saturation ratings for each subject across trials for stabilized stimuli; triangles are for unstabilized stimuli. Error bars represent +/-1 standard error of the mean across trials. ** indicates that there was a main effect of intensity where p < 0.01. **b)** Proportion of trials that subjects described stimuli of each intensity as green, as opposed to red or white. Circles are green proportions for stabilized stimuli; triangles are for unstabilized stimuli. ** indicates that there was a main effect of intensity where p < 0.01. In both panels, data are collapsed across size conditions, and horizontally offset from the true abscissa value for aesthetic purposes.

In Figure 3, both eye motion conditions are combined to examine the joint effects of stimulus size and intensity. Since previous studies have shown that increasing intensity does not significantly change hue or saturation perception for single cone stimuli, in the current study we only presented such stimuli at threshold intensity. The nature of the data precludes a straightforward statistical assessment of an interaction between size and intensity. However, the difference in the apparent slopes of the functions shown in Figure 3 can indicate the existence of interactions in the independent variables. Insofar as that is the case, there appears to be such an interaction on the proportion of green responses, and potentially a milder interaction on mean saturations.

**Figure 3:**
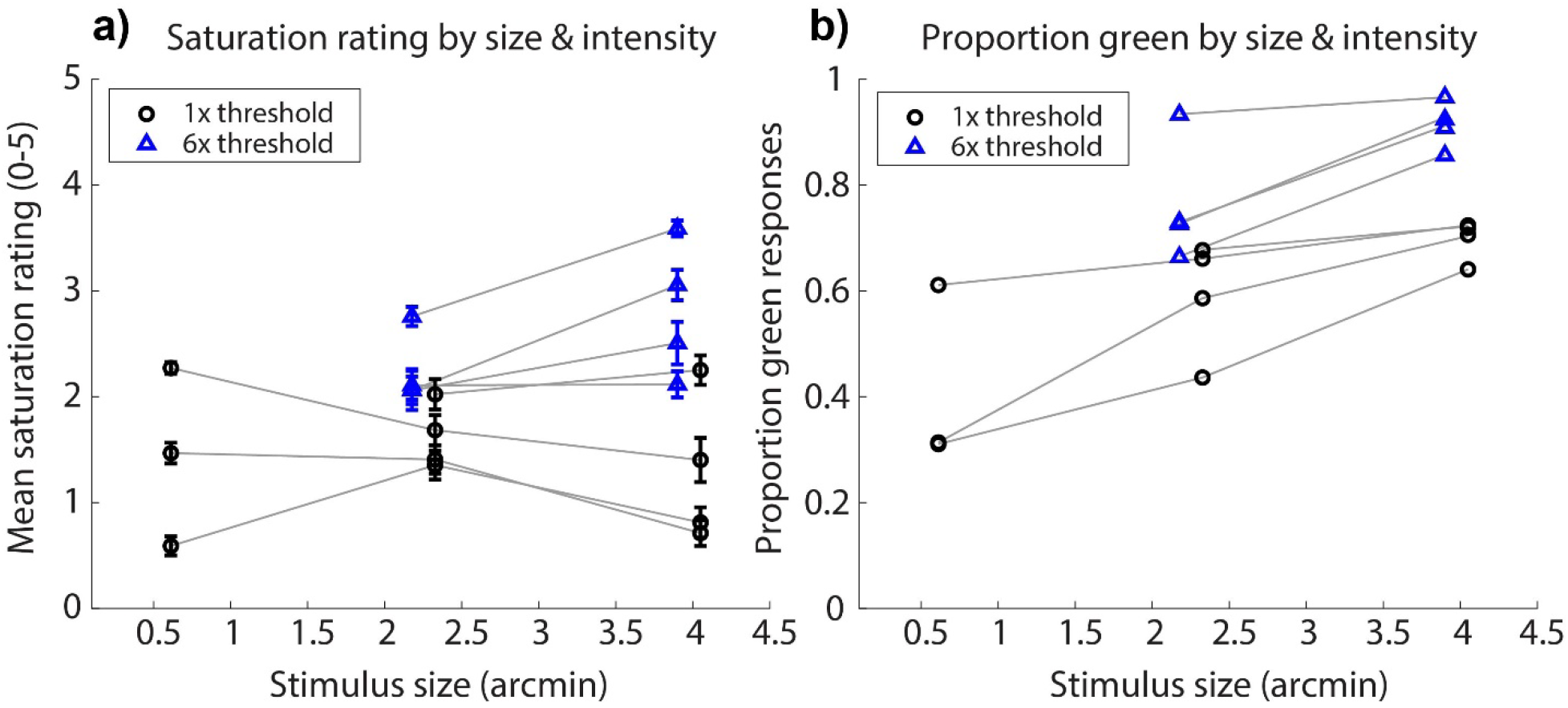
**a)** Saturation rating for each stimulus size condition. Circles are mean saturation ratings for each subject across trials for 1x threshold stimuli; triangles are for 6x threshold stimuli. Error bars represent +/-1 standard error of the mean across trials. **b)** Proportion of trials that subjects described stimuli of each size as green, as opposed to red or white. Circles are green proportions for 1x threshold stimuli; triangles are for 6x threshold stimuli. In both panels, data are collapsed across eye motion conditions, and horizontally offset from the true abscissa value for aesthetic purposes.

One of the main effective differences between the eye motion conditions is that unstabilized stimuli are likely to traverse a greater distance across the retina, and thus stimulate more cones than stabilized stimuli. While there was no main effect of eye motion condition in any comparison, we hypothesized that trials in which the stimulus stimulated more cones might lead to higher saturation rankings and/or increased likelihoods of reporting that the stimulus appeared green. To investigate this, we first sorted trials by stimulus size, as different stimulus sizes result in different distributions of stimulated cones. We then sorted trials into bins based on the number of cones stimulated over the course of each trial. The process to count the number of stimulated cones per trial is as follows: First, the stimulus size and exact trajectory of each stimulus for each trial is known. Counting the number of cones simply involved counting the total number of cones that engaged at any point with the stimulus over the course of each trial. Figure 4 shows two examples, where 11 cones are engaged for a stabilized stimulus and 25 cones are engaged for the same sized stimulus that is naturally moving. The associated movie in the **Supplementary Material** shows the manner in which all stimulated cones are engaged over 15 frames. Note that even in the ‘stabilized’ trials, there is residual motion and so there is some variability in number of cones stimulated for those trials also. For each bin, we averaged across all trials for all subjects to compute a mean saturation and computed the proportions of each type of hue response. The results of this analysis are shown in Figure 5.

**Figure 4:**
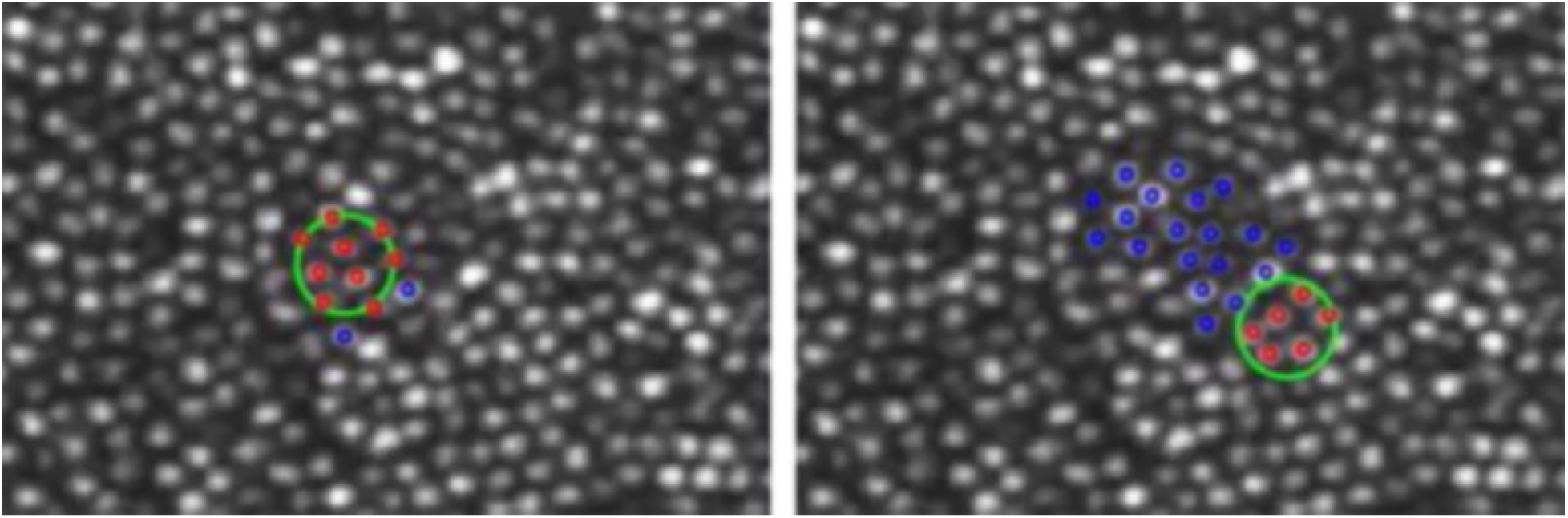
An AOSLO image of the cone mosaic of S10003, averaged across many frames. This is the final frame of video that is included in the **Supplementary Material**. The gray spots are cone photoreceptors. The large green circle and small red and blue circles were added using image editing software and are based on the known size, position, and trajectory of the stimulus for each experimental trial. The green circle represents the location and approximate size of the 543-nm spot of light; on these trials, the stimulus was 2.3-arcmin in size (not accounting for light spread due to diffraction). The red circles show the centers of the cones which are being stimulated in the current frame and the blue circles indicate cones that were stimulated in previous frames. The left frame is from a stabilized condition. Stabilization errors cause more cones to be stimulated than what falls under the stimulus on any given frame. The right frame is from an unstabilized condition. In this case the eye motion results in stimulation of many more cones on a typical trial.

**Figure 5:**
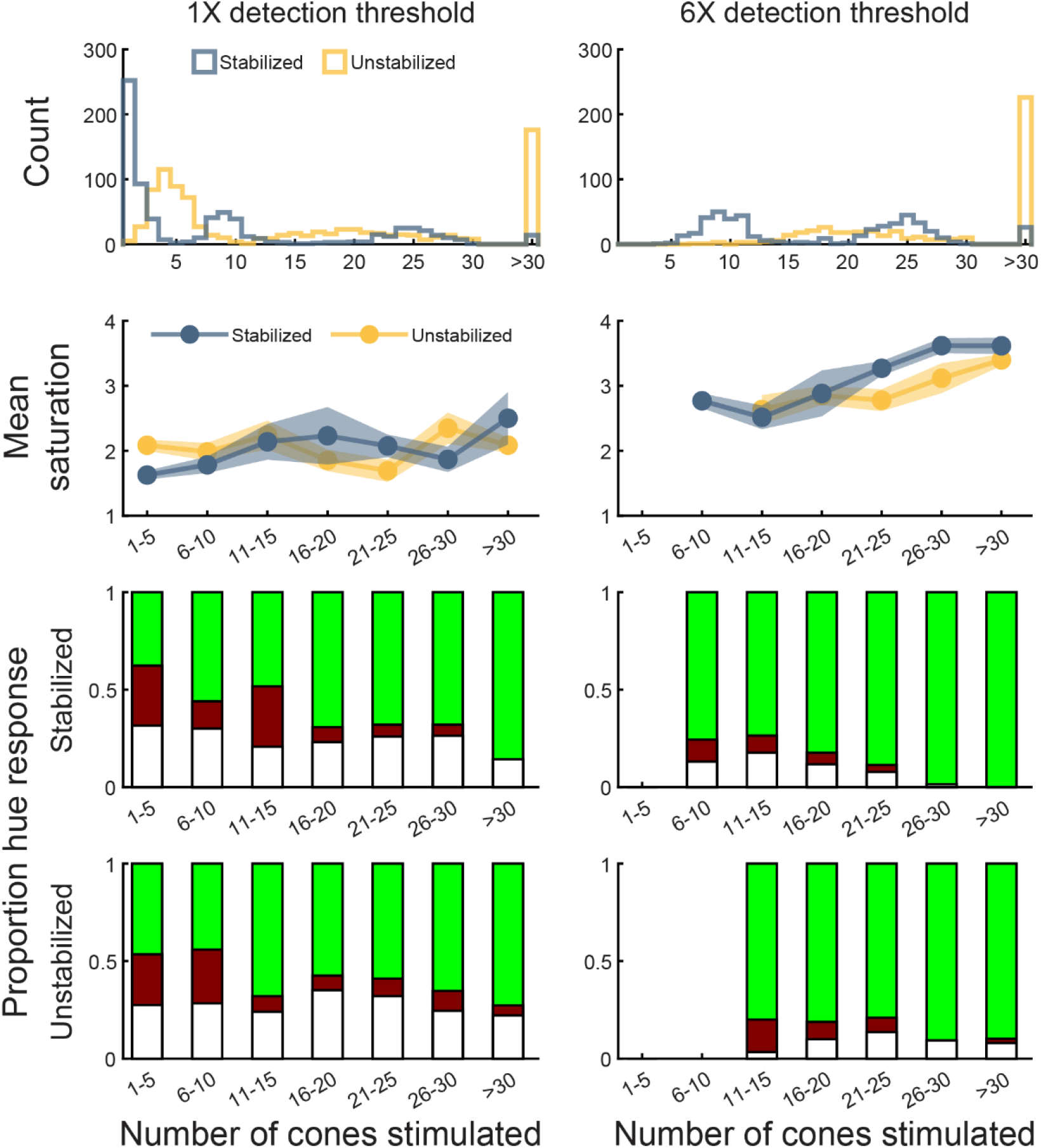
Hue and saturation as a function of cones stimulated. Data were separated by intensity condition (left column: 1X threshold; right column: 6X threshold). Top row: histograms depicting the number of cones stimulated per trial for stabilized (blue) and unstabilized (yellow) stimuli. Data from all stimulus sizes are shown. Second row: mean saturation ratings for stabilized (blue circles) and unstabilized (yellow circles). The shaded regions delimit ±1 standard error of the mean for each retinal motion condition. Third row: hue responses as a function of cones stimulated by retinally stabilized stimuli. The shaded green, red, and white regions represent the proportion of trials rated with those terms. Fourth row: proportional hue responses for unstabilized stimuli. In all panels, data were pooled across subjects. In the second through fourth panels, bins containing fewer than 10 trials (∼0.4% of all analyzed trials) were excluded to more clearly represent summary trends.

The top row of Figure 5 shows histograms for number of cones stimulated per trial for the low (left column) and high (right column) intensity conditions. Data from each stimulus size are included in this analysis. For the low intensity condition, stabilized trials (blue) reveal three distinct clusters that correspond to the three stimulus sizes used in the study. In the high intensity condition (right), the smallest stimulus size was not used, hence only two distributions are apparent. By comparison, unstabilized trials (yellow) exhibit a higher median and greater variation in number of cones encountered, which reflects both expected eye motion behaviors and the exclusion criteria used. In the remaining panels of Figure 5, bins containing fewer than 10 trials (∼0.4% of all trials) were excluded to illustrate the summary trends more clearly.

The second row of Figure 5 shows mean saturation ratings as a function of cones stimulated for both stabilized (blue) and unstabilized (yellow) stimuli. Considered solely in terms of the number of cones engaged by the stimulus, there is no obvious difference in saturation ratings for the two retinal motion conditions at either low or high intensity. The third row shows the proportion of stabilized trials rated as green, red or white as a function of number of cones stimulated. The fourth row shows the same results for unstabilized stimuli. In these plots, the relative size of each colored portion represents the proportion of that hue response. (e.g., if the bar is half green, then half the trials for that bin resulted in green percepts). Consistent with the results shown in Figures 1 and 3, increasing both size and stimulus intensity increases the likelihood that our 543 nm stimulus will be correctly judged as green. For the high-intensity, stabilized trials (third row, right column), our data suggest that stimulating a half-dozen or more cones is sufficient to trigger a green sensation on a majority of trials. Comparing across retinal motion conditions, there does not appear to be any significant difference in hue naming trends.

## Discussion

Here we summarize and explain our results and put them into the context of previous literature where appropriate. Numerous studies have examined color appearance with multiple factors including stimulus size, duration, retinal location, intensity and background illumination. Aside from a handful of studies, none of them used tiny, cone-sized stimuli achieved with adaptive optics or used retinal stabilization. Rather, the focus of most previous studies was on the appearance of larger spots, which elicited near-veridical percepts, and particular emphasis was placed on comparisons between fovea and larger eccentricities than were tested in our study. As such, extrapolating conditions from previous work to those of the current study needs to be done with caution.

### Size & Intensity

In the present study, test spots ranged in diameter from half an arcminute to four arcminutes, stimulating on average for any given frame from one to nineteen cones, respectively. Varying the spot size of 543-nm light over this range had no significant impact on mean saturation ratings at detection threshold but did change the likelihood that subjects reported that the stimulus looked green, rather than red or white (Figure 1). Stimuli presented at six times detection threshold were rated as being significantly more saturated and were significantly more likely to be described as green than stimuli presented at detection threshold (Figure 2). In addition to the main effect of light intensity on our dependent variables, the data suggest an interaction between stimulus size and intensity such that higher intensities were more effective at eliciting greener and more saturated percepts when stimuli were larger (Figure 3).

Previous studies have shown that when cones are stimulated individually, the resulting hue percepts depend on the spectral identities of the cones rather than the spectral composition of the stimulus (Hofer et al., 2005; Sabesan et al., 2016; Schmidt et al., 2018). The restricted spatial profile of a single-cone flash prevents the brain from directly measuring how the other cone types would respond to the stimulus. As a result, the color sensations elicited under these conditions are often not veridical, underscoring how the inference processes that facilitate trichromatic reconstruction are constrained by the quality of the information available to them (Brainard, 2015; Brainard, Williams, & Hofer, 2008; Zhang, Cottaris, & Brainard, 2022). Schmidt et al. (2019) extended these methods to study color appearance when pairs of cones were stimulated with 543-nm light, demonstrating that, compared to single-cone stimulation, pairs of like-type cones gave more saturated percepts – sensations which were still linked to cone type and thus sometimes non-veridical. By contrast, different-type (i.e., L/M) cone pairs produced largely achromatic (i.e., desaturated) sensations. These results suggest that stimulating two cones – even if those cones belong to different spectral classes – is not sufficient to guarantee veridical color perception. Using the same Bayesian framework that had been successfully applied to model the highly variable hue sensations evoked by single-cone flashes, Brainard et al. (2008) found that the mosaic response to an 8 arcmin, 550-nm spot contained enough information for the stimulus to be consistently reconstructed as greenish. Taken together, these results place bounds on the spatial scale at which reliable hue signals are generated and veridical color perception is likely to emerge. This prediction about the critical area for color appearance is consistent with our data, which showed that engaging 6 or more cones with a suprathreshold spot of 543-nm light ensured it would be judged as green a majority of the time (**Figure 5**).

While increasing the stimulus diameter to encompass a half-dozen or more cones increased the likelihood that 543-nm would be judged as green, the effect of stimulus size on apparent saturation was less straightforward and appeared to depend on stimulus intensity: saturation ratings were invariant to spot diameter for intensities near detection threshold, while suprathreshold spots appeared more saturated as stimulus size was increased, although this trend was not significant (**Figure 3a**). When delivered over relatively large fields (> 1°), near-monochromatic wavelengths like those used in our study are known to elicit the most saturated percepts (Onley, Klingberg, Dainoff, & Rollman, 1963). However, Gordon and Abramov (1977) showed that decreasing stimulus size from 1.5° to 5 arcmin reduced the perceived saturation of foveally-viewed suprathreshold (1200 td) spectral lights, with the most prominent reduction occurring near our 543-nm test wavelength. Our measurements using 2.3 and 4.0-arcmin suprathreshold flashes suggest the size dependency of saturation demonstrated by Gordon and Abramov (1977) extends to even smaller stimulus sizes.

Although the saturation ratings of our suprathreshold spots followed previously reported trends, the saturations associated with our near-threshold stimuli were invariant to stimulus size (**Figure 3a**). How can we explain this finding? It has been suggested that perceived saturation depends on the ratio of activity in opponent and non-opponent neural channels (Guth, 1991; Jameson & Hurvich, 1955). Spectrally-opponent, parvocellular-projecting midget retinal ganglion cells (RGCs) are thought to be the primary contributors to the red-green dimension of color vision (Dacey, 2000), whereas parasol RGCs, which sum L and M cone activity in a non-opponent fashion and project to the magnocellular pathway, have been proposed as a putative neural substrate for an achromatic visual channel (Lee, Martin, & Valberg, 1988; Lee, Pokorny, Smith, Martin, & Valberg, 1990) but see Lennie and D’Zmura (1998) and Lennie, Pokorny, and Smith (1993).

The saturation invariance we observed at detection threshold implies that the ratio of activity in the chromatic and achromatic channels was approximately constant across the range of sizes and intensities used. The intensities used in the current study were defined by the detection thresholds for each stimulus size, which in turn were presumably determined by the combined activity of the mechanisms engaged by each spot (Chaparro, Stromeyer, Kronauer, & Eskew, 1994; Cole, Hine, & McIlhagga, 1993). In the peripheral retina, single-cone modulations elicit similar magnitudes of spiking activity in midget and parasol RGCs (P. H. Li et al., 2014). In the case of medium and large stimuli, which encompass a target cone and either one or two concentric rings of surrounding cones, respectively, parasol cells are likely to be the most sensitive RGC type because they receive input from multiple cones. Thus, the medium and large stimulus threshold intensities may have been below threshold for an individual midget cell, which likely draw their excitatory input from single cones at our test eccentricity. Considering the properties of individual RGCs, one might conclude that the balance of excitation for larger stimulus sizes would shift to the achromatic mechanism, and perceived saturation at detection threshold would therefore decrease as spot size increases. It is important to note, however, that the relatively low spatial resolution for veridical perception of red-green equiluminant gratings implies that chromatic signals from individual midget RGCs are pooled at a post-retinal site (Neitz et al., 2020; Sekiguchi, Williams, & Brainard, 1993). As spot diameter was increased, it is conceivable that spatial summation of subthreshold chromatic signals preserved the balance of activity in the post-receptoral mechanisms, in turn ensuring that perceived saturation remained stable across the narrow range of stimulus sizes employed in this study.

Our data also demonstrate that, when these larger stimuli were presented many times above threshold, saturation ratings were slightly higher and exhibited a clearer dependency on stimulus size. These results suggest that, as overall stimulus energy is increased, the balance of excitation may begin to shift to the parvocellular system. The bias towards chromatic pathways in the suprathreshold regime may relate to the observation that the responses of magnocellular neurons tend to saturate at much lower contrasts than their parvocellular counterparts (Kaplan & Shapley, 1986). While we have attempted to interpret our results on small-spot hue and saturation perception within a framework built around well-known properties (i.e., spectral sampling, spatial pooling, contrast gain) of early visual neurons, we also acknowledge that, as evidenced by recent electrophysiological studies, color signals are subjected to a diverse and complex set of transformations as they enter the cortex and traverse the visual hierarchy (De Valois, Cottaris, Elfar, Mahon, & Wilson, 2000; Horwitz & Hass, 2012; M. Li et al., 2022; P. Li, Garg, Zhang, Rashid, & Callaway, 2022; Liu et al., 2020). The comparatively limited nature of our dataset makes it difficult to draw firm links between our perceptual results and these higher-level processing stages.

Finally, we consider the impact test eccentricity has on the relationship between stimulus size and perceived saturation. In general, patches of light of fixed size appear less saturated when test eccentricity is increased (Mogi, Sakurai, Ishikawa, & Ayama, 2021; Weitzman & Kinney, 1969). However, the effect of eccentricity on perceived saturation is likely more complicated at the retinal locations used in this study (0.5 – 1 degree from the fovea), particularly when small spots are used. Ingling, Scheibner, and Boynton (1970) found that small (3 arcmin), middle-wavelength flashes appeared, on average, slightly more saturated when delivered at 1 deg eccentricity compared to when they were viewed foveally. These results may be a consequence of the tritanopic nature of the foveola, which is devoid of S-cone receptors within the central 20 arcmin (Curcio et al., 1991; Williams, MacLeod, & Hayhoe, 1981). While experimental data on saturation perception in congenital tritanopia is scant due to its extremely low incidence, theoretical modeling suggests that the absence of an S-cone signal would reduce the apparent saturation of middle-wavelength lights like those used in this study (Hurvich & Jameson, 1955). By comparison, S-cone density is higher at our test eccentricity, and although these receptors still only make up on a small fraction (5 to 7%) of the overall cone population (Curcio et al., 1991), the likelihood that our small test flashes would be sampled by all three cone types is substantially higher in the parafovea. Repeating the present study at the foveal center may lead to a slightly larger estimate of the critical area for reliable hue and saturation perception of mid-spectrum lights.

### Motion

Previous research on color perception used single-cone, stabilized stimuli on the retina so that the same cones or pairs of cones could be repeatedly stimulated over the course of a trial. While monochromatic 543-nm light viewed under more normal conditions (e.g., shone onto a white surface) looks invariably green, this same light stabilized on single cones may appear as a variety of hues. The central nervous system integrates information over space and time, and stabilization may effectively reduce the amount of spatiotemporal information available from which to build a color percept. When the motion of a larger chromatic stimulus on the retina is minimized experimentally, the stimulated region tends to take on the color appearance of the surrounding field as it fades from view (Krauskopf, 1963; Larimer & Piantanida, 1988). When the salience of the perceptual boundary between the target and background is increased – either by sharpening the chromatic transition between background and target or by introducing a small amount of target motion – the filling-in effect is reduced (Friedman, Zhou, & von der Heydt, 1999). While the aforementioned studies used larger (>1 deg) stimuli to reveal phenomena that occur over relatively long (>3 s) time courses, we nonetheless hypothesized that allowing our small, brief flashes to move freely across the retina during the subject’s natural eye motion may increase the likelihood of a veridical percept (green, in this case). Within the range of parameters studied here, this does not appear to be the case. We saw no significant effect of eye motion condition in any statistical comparison, nor was the number of cones traversed over the course of a trial predictive of saturation rating or hue name given to a stimulus. There is little question that increasing the number of cones stimulated simultaneously leads to a higher frequency of green percepts, but it appears that at these time scales (0.5 sec stimulus duration) information from sequential cone excitations is not being integrated in service to hue perception. By comparison, eye motion is leveraged to improve other tasks; for instance, fixational eye movements have been shown to increase contrast sensitivity (Rucci et al., 2007) and improve visual acuity for high spatial frequency stimuli (Anderson et al., 2020; Ratnam et al., 2017).

The lack of a significant effect of eye motion in the current study may reflect either a difference in the size and duration of the stimulus that are needed for benefits in hue perception to occur, or a more fundamental difference between chromatic and luminance processing. Given that in the unstabilized condition, stimuli were initially delivered to the same target locations as in the stabilized condition, the first cone or cones stimulated may be essential for determining the final percept at these time scales. Longer presentation durations for unstabilized stimuli could result in more saturated or greener percepts.

### Color appearance in relation to the cone mosaic

Stimulating single cones and pairs of cones with spots of 543-nm light results in a variety of hue percepts, which can include yellowish and blueish components but vary mainly along a red-green axis in color-opponent space (Schmidt et al., 2018). We replicated these findings and determined that this overall range of percepts persists at larger spatial scales. Larger stimuli were more likely to be described as green than smaller stimuli, but even the largest spot (which had ∼54 times the area of the smallest spot) was sometimes described as white or red. We observed some individual variation in response patterns. Notably, S10003 was the most likely to describe stimuli as red overall, while S20210 rarely used this hue name, even for the smallest stimuli. For other research purposes, the topography of the S, M and L-cones and corresponding L:M ratios of these subjects were determined optoretinographically using an AO optical coherence tomography system (Pandiyan, Bertelli, et al., 2020; Pandiyan, Jiang, et al., 2020), at similar eccentricities but for a different patch of cones than were used for this experiment. S10003 was found to have an L:M ratio of 1.35, while S20210 had an L:M ratio of 0.84. While these are only two subjects, this observation suggests that even for stimuli that encompass the receptive fields of multiple ganglion cells, the observed statistical relationship between cone spectral identity and hue percept may endure.

## Conclusions

This study examined the effects of stimulus size, intensity, and retinal stabilization on saturation and hue perception for very small spots of monochromatic 543-nm light. Making the spots larger and brighter elicited more frequent perceptions of green, which is the veridical appearance of this light (i.e., 543-nm light looks green when freely viewed). Increasing the intensity, but not the size, of the spots led to more saturated percepts overall. The data suggest an interaction between size and intensity that can possibly be explained by the known properties of retinal ganglion cells. Whether the stimulus was stabilized on the retina, as in previous studies, or allowed to move across the retina made little difference to hue or saturation perception, either overall or when considered by the number of cones stimulated sequentially per trial. The asynchronous excitations of contiguous groups of cones over the timescales measured here do not appear to be a useful source of information about hue or saturation.

## Supporting information

Movie related to figure 4

## Acknowledgments

The authors thank Professor Ramkumar Sabesan and Vimal Pandiyan for their time and effort in providing AO-OCT measurements of L:M ratios, and to Pavan Tiruveedhula for technical assistance.

## Supplementary Material

15 frame movie indicating the cones that engage with the stimulus over the course of a single trial. The left frame shows a stabilized sequence and the right frame shows a sequence with natural eye motion.

## Notes

<u>**Funding:**</u> This research was supported by the following grants:

### Competing Interest Statement

Disclosures:
JEV: none
AEB: none
WST: Patent assigned to the University of California
AR: Patent assigned to the University of California; Patent assigned to the University of Houston & the University of Rochester

## References

Anderson, A. G., Ratnam, K., Roorda, A., & Olshausen, B. A. (2020). High-acuity vision from retinal image motion. Journal of Vision, 20(7), 34–34. doi:10.1167/jov.20.7.34

Arathorn, D. W., Yang, Q., Vogel, C. R., Zhang, Y., Tiruveedhula, P., & Roorda, A. (2007). Retinally Stabilized Cone-Targeted Stimulus Delivery Optics Express, 15, 13731–13744.

Atchison, D. A., & Smith, G. (2005). Chromatic dispersions of the ocular media of human eyes. Journal of the Optical Society of America A, 22(1), 29–37.

Brainard, D. H. (1997). The Psychophysics Toolbox. Spatial Vision, 10(4), 433–436.

Brainard, D. H. (2015). Color and the Cone Mosaic. Annual Reviews Vision Science, 1, 519–546. doi:10.1146/annurev-vision-082114-035341

Brainard, D. H., Williams, D. R., & Hofer, H. (2008). Trichromatic reconstruction from the interleaved cone mosaic: Bayesian model and the color appearance of small spots. Journal of Vision, 8(5).

Chaparro, A., Stromeyer, C. F., 3rd, Kronauer, R. E., & Eskew, R. T., Jr. (1994). Separable redgreen and luminance detectors for small flashes. Vision Research, 34(6), 751–762. doi:10.1016/0042-6989(94)90214-3

Cole, G. R., Hine, T., & McIlhagga, W. (1993). Detection mechanisms in L-, M-, and S-cone contrast space. Journal of the Optical Society of America A, 10(1), 38–51. doi:10.1364/josaa.10.000038

Curcio, C. A., Allen, K. A., Sloan, K. R., Lerea, C. L., Hurley, J. B., Block, I. B., & Milam, A. H. (1991). Distribution and morphology of human cone photoreceptors stained with anti-blue opsin. Journal of Comparitive Neurology, 312, 610–624.

Curcio, C. A., Sloan, K. R., Kalina, R. E., & Hendrickson, A. E. (1990). Human photoreceptor topography. Journal of Comparitive Neurology, 292, 497–523.

Dacey, D. M. (2000). Parallel pathways for spectral coding in primate retina. Annual Reviews Neuroscience, 23, 743–775. doi:10.1146/annurev.neuro.23.1.743

De Valois, R. L., Cottaris, N. P., Elfar, S. D., Mahon, L. E., & Wilson, J. A. (2000). Some transformations of color information from lateral geniculate nucleus to striate cortex. Proceedings of the National Academy of Sciences, 97(9), 4997–5002.

Domdei, N., Linden, M., Reiniger, J. L., Holz, F. G., & Harmening, W. M. (2019). Eye tracking-based estimation and compensation of chromatic offsets for multi-wavelength retinal microstimulation with foveal cone precision. Biomedical Optics Express, 10(8), 4126–4141. doi:10.1364/BOE.10.004126

Friedman, H. S., Zhou, H., & von der Heydt, R. (1999). Color filling-in under steady fixation: behavioral demonstration in monkeys and humans. Perception, 28(11), 1383–1395. doi:10.1068/p2831

Gordon, J., & Abramov, I. (1977). Color vision in the peripheral retina. II. Hue and saturation. Journal of the Optical Society of America, 67(2), 202–207. doi:10.1364/josa.67.000202

Grieve, K., Tiruveedhula, P., Zhang, Y., & Roorda, A. (2006). Multi-wavelength imaging with the adaptive optics scanning laser ophthalmoscope. Optics Express, 14(25), 12230–12242.

Guth, S. L. (1991). Model for color vision and light adaptation. Journal of the Optical Society of America A, 8(6), 976–993. doi:10.1364/josaa.8.000976

Harmening, W. M., Tiruveedhula, P., Roorda, A., & Sincich, L. C. (2012). Measurement and correction of transverse chromatic offsets for multi-wavelength retinal microscopy in the living eye. Biomedical Optics Express, 3(9), 2066–2077.

Harmening, W. M., Tuten, W. S., Roorda, A., & Sincich, L. C. (2014). Mapping the perceptual grain of the human retina. Journal of Neuroscience, 34(16), 5667–5677.

Hofer, H., Singer, B., & Williams, D. R. (2005). Different sensations from cones with the same pigment. Journal of Vision, 5(5), 444–454.

Horwitz, G. D., & Hass, C. A. (2012). Nonlinear analysis of macaque V1 color tuning reveals cardinal directions for cortical color processing. Nature Neuroscience, 15(6), 913–919. doi:10.1038/nn.3105

Hurvich, L. M., & Jameson, D. (1955). Some quantitative aspects of an opponent-colors theory. II. Brightness, saturation, and hue in normal and dichromatic vision. Journal of the Optical Society of America, 45(8), 602–616. doi:10.1364/josa.45.000602

Ingling, C. R., Jr., Scheibner, H. M., & Boynton, R. M. (1970). Color naming of small foveal fields. Vision Research, 10(6), 501–511. doi:10.1016/0042-6989(70)90006-4

Jameson, D., & Hurvich, L. M. (1955). Some quantitative aspects of an opponent-colors theory. I. Chromatic responses and spectral saturation. Journal of the Optical Society of America, 45(7), 546–552.

Kaplan, E., & Shapley, R. M. (1986). The primate retina contains two types of ganglion cells, with high and low contrast sensitivity. Proc Natl Acad Sci U S A, 83(8), 2755–2757. doi:10.1073/pnas.83.8.2755

Kleiner, M., Brainard, D., Pelli, D., Ingling, A., Murray, R., & Broussard, C. (2007). What’s new in Psychtoolbox-3? Perception, 36(14), 16.

Krauskopf, J. (1963). Effect of retinal image stabilization on the appearance of heterochromatic targets. Journal of the Optical Society of America, 53, 741–744. doi:10.1364/josa.53.000741

Krauskopf, J., & Srebro, R. (1965). Spectral sensitivity of color mechanisms: derivation from fluctuations of color appearance near threshold. Science, 150(3702), 1477–1479. doi:10.1126/science.150.3702.1477

Kruskal, W. H., & Wallis, W. A. (1952). Use of ranks in one-criterion variance analysis. Journal of the American Statistical Association, 47(260), 583–621.

Larimer, J., & Piantanida, T. (1988). The impact of boundaries on color: Stabilized image studies. SPIE conference on Image Processing, Analysis, Measurement, and Quality, 901, 241–247.

Lee, B. B., Martin, P. R., & Valberg, A. (1988). The physiological basis of heterochromatic flicker photometry demonstrated in the ganglion cells of the macaque retina. Journal of Physiology, 404, 323–347. doi:10.1113/jphysiol.1988.sp017292

Lee, B. B., Pokorny, J., Smith, V. C., Martin, P. R., & Valberg, A. (1990). Luminance and chromatic modulation sensitivity of macaque ganglion cells and human observers. Journal of the Optical Society of America A, 7(12), 2223–2236. doi:10.1364/josaa.7.002223

Leibowitz, H. (1954). The use and calibration of the Maxwellian view in visual instrumentation. American Journal of Psychology, 67(3), 530–532.

Lennie, P., & D’Zmura, M. (1998). Mechanisms of color vision. CRC Critical Reviews in Neurobiology, 3(4), 333–400.

Lennie, P., Pokorny, J., & Smith, V. C. (1993). Luminance. Journal of the Optical Society of America A, 10(6), 1283–1293.

Li, M., Ju, N., Jiang, R., Liu, F., Jiang, H., Macknik, S., … Tang, S. (2022). Perceptual hue, lightness, and chroma are represented in a multidimensional functional anatomical map in macaque V1. Progress in Neurobiology, 212, 102251.

Li, P., Garg, A. K., Zhang, L. A., Rashid, M. S., & Callaway, E. M. (2022). Cone Opponent Functional Domains in Primary Visual Cortex Combine Signals for Color Appearance Mechanisms. Nat Commun, 13(1), 19.

Li, P. H., Field, G. D., Greschner, M., Ahn, D., Gunning, D. E., Mathieson, K., … Chichilnisky, E. J. (2014). Retinal representation of the elementary visual signal. Neuron, 81(1), 130–139.

Liu, Y., Li, M., Zhang, X., Lu, Y., Gong, H., Yin, J., … Wang, W. (2020). Hierarchical Representation for Chromatic Processing across Macaque V1, V2, and V4. Neuron, 108(3), 538–550 e535. doi:10.1016/j.neuron.2020.07.037

Mogi, S., Sakurai, M., Ishikawa, T., & Ayama, M. (2021). Color appearance of small stimuli presented in central and near peripheral visual fields. Color Research and Application, 46(4), 722–739. doi:10.1002/col.22610

Mozaffari, S., LaRocca, F., Jaedicke, V., Tiruveedhula, P., & Roorda, A. (2020). Wide-vergence, multi-spectral adaptive optics scanning laser ophthalmoscope with diffraction-limited illumination and collection. Biomedical Optics Express, 11(3), 1617–1632. doi:10.1364/BOE.384229

Neitz, A., Jiang, X., Kuchenbecker, J. A., Domdei, N., Harmening, W., Yan, H., … Sabesan, R. (2020). Effect of cone spectral topography on chromatic detection sensitivity. Journal of the Optical Society of America A, 37(4), A244–A254. doi:10.1364/JOSAA.382384

Onley, J. W., Klingberg, C. L., Dainoff, M. J., & Rollman, G. B. (1963). Quantitative Estimates of Saturation. Journal of the Optical Society of America, 53(4), 487–493. doi:10.1364/JOSA.53.000487

Pandiyan, V. P., Bertelli, A. M., Kuchenbecker, J. A., Boyle, K. C., Ling, T., Chen, Z. C., … Sabesan, R. (2020). The optoretinogram reveals the primary steps of phototransduction in the living human eye. Sci Adv, 6, eabc1124. doi:doi.org/10.1126/sciadv.abc1124

Pandiyan, V. P., Jiang, X., Maloney-Bertelli, A., Kuchenbecker, J. A., Sharma, U., & Sabesan, R. (2020). High-speed adaptive optics line-scan OCT for cellular-resolution optoretinography. Biomedical Optics Express, 11(9), 5274–5296. doi:https://doi.org/10.1364/BOE.399034

Pelli, D. G. (1997). The VideoToolbox software for visual psychophysics: transforming numbers into movies. Spatial Vision, 10(4), 437–442.

Ratnam, K., Domdei, N., Harmening, W. M., & Roorda, A. (2017). Benefits of retinal image motion at the limits of spatial vision. Journal of Vision, 17(1), 30. doi:10.1167/17.1.30

Roorda, A., Romero-Borja, F., Donnelly, W. J., Queener, H., Hebert, T. J., & Campbell, M. C. W. (2002). Adaptive optics scanning laser ophthalmoscopy. Optics Express, 10(9), 405–412.

Rucci, M., Iovin, R., Poletti, M., & Santini, F. (2007). Miniature eye movements enhance fine spatial detail. Nature, 447(7146), 852–855.

Rushton, W. A. (1972). Pigments and signals in colour vision. Journal of Physiology, 220(3), 1–31P. doi:10.1113/jphysiol.1972.sp009719

Sabesan, R., Schmidt, B. P., Tuten, W. S., & Roorda, A. (2016). The elementary representation of spatial and color vision in the human retina. Sci Adv, 2(9), e1600797. doi:10.1126/sciadv.1600797

Schmidt, B. P., Boehm, A. E., Foote, K. G., & Roorda, A. (2018). The spectral identity of foveal cones is preserved in hue perception. Journal of Vision, 18(11), 19. doi:10.1167/18.11.19

Schmidt, B. P., Boehm, A. E., Tuten, W. S., & Roorda, A. (2019). Spatial summation of individual cones in human color vision. PLoS One, 14(7), e0211397. doi:10.1371/journal.pone.0211397

Sekiguchi, N., Williams, D. R., & Brainard, D. H. (1993). Efficiency in detection of isoluminant and isochromatic interference fringes. Journal of the Optical Society of America A, 10, 2118–2133.

Wang, Y., Bensaid, N., Tiruveedhula, P., Ma, J., Ravikumar, S., & Roorda, A. (2019). Human foveal cone photoreceptor topography and its dependence on eye length. Elife, 8. doi:10.7554/eLife.47148

Watson, A. B., & Pelli, D. G. (1983). QUEST: a Bayesian adaptive psychometric method. Perceptual Psychophysics, 33(2), 113–120.

Weitzman, D. O., & Kinney, J. A. S. (1969). Effect of Stimulus Size, Duration, and Retinal Location upon the Appearance of Color. Journal of the Optical Society of America, 59(5), 640–640. doi:10.1364/JOSA.59.000640

Williams, D. R., MacLeod, D. I. A., & Hayhoe, M. M. (1981). Foveal tritanopia. Vision Research, 21, 1341–1356.

Yang, Q., Arathorn, D. W., Tiruveedhula, P., Vogel, C. R., & Roorda, A. (2010). Design of an integrated hardware interface for AOSLO image capture and cone-targeted stimulus delivery. Optics Express, 18(17), 17841–17858.

Zhang, L. Q., Cottaris, N. P., & Brainard, D. H. (2022). An image reconstruction framework for characterizing initial visual encoding. Elife, 11. doi:10.7554/eLife.71132

